# Hybrid dysgenesis in *Drosophila simulans* associated with a rapid invasion of the *P*-element

**DOI:** 10.1101/031781

**Authors:** Tom Hill, Christian Schlötterer, Andrea J. Betancourt

## Abstract

In a classic example of the invasion of a species by a selfish genetic element, the *P-*element was horizontally transferred from a distantly related species into *Drosophila melanogaster*. Despite causing ‘hybrid dysgenesis’, a syndrome of abnormal phenotypes that include sterility, the *P-*element spread globally in the course of a few decades in *D. melanogaster*. Until recently, its sister species, including *D. simulans*, remained *P-*element free. Here, we find a hybrid dysgenesis-like phenotype in the offspring of crosses between *D. simulans* strains collected in different years; a survey of 181 strains shows that around 20% of strains induce hybrid dysgenesis. Using genomic and transcriptomic data, we show that this dysgenesis-inducing phenotype is associated with the invasion of the *P-*element. To characterize this invasion temporally and geographically, we survey 631 *D. simulans* strains collected on three continents and over 27 years for the presence of the *P-*element. We find that the *D. simulans P-*element invasion occurred rapidly and nearly simultaneously in the regions surveyed, with strains containing *P-*elements being rare in 2006 and common by 2014. Importantly, as evidenced by their resistance to the hybrid dysgenesis phenotype, strains collected from the latter phase of this invasion have adapted to suppress the worst effects of the *P-*element.

**Author Summary:** Some genes perform necessary organismal functions, others hijack the cellular machinery to replicate themselves, potentially harming the host in the process. These ‘selfish genes’ can spread through genomes and species; as a result, eukaryotic genomes are typically saddled with large amounts of parasitic DNA. Here, we chronicle the surprisingly rapid global spread of a selfish transposable element through a close relative of the genetic model, *Drosophila melanogaster*. We see that, as it spreads, the transposable element is associated with damaging effects, including sterility, but that the flies quickly adapt to the negative consequences of the transposable element.

## Introduction

Discovered in the 1950’s by Barbara McClintock [1], transposable elements (TEs) are the ‘ultimate parasite’ [2, 3]. They are genetically simple, with autonomous elements consisting of as little as one protein coding gene, but remarkably successful, comprising much of the bulk of large eukaryotic genomes [4, 5]. Those that persist do so by inserting copies of themselves into new genomic locations faster than they are eliminated. TEs can also be a source of beneficial mutations; particular TE insertions have been implicated in a number of adaptive phenotypes (reviewed in [6]), and some TE families now form essential components of genomes [7, 8]. But most TE insertions are probably deleterious, as insertions can damage genes and may impose other fitness costs on their host [9–11]. In fact, the mutagenic properties of transposons have been harnessed by molecular geneticists to disrupt thousands of *Drosophila* genes (*e.g.*, [12]).

In the short-term, transposable elements are vertically transmitted, and so can only cause limited damage to their hosts without extinguishing themselves. In the long-term, however, they are also horizontally transmitted [13–15], and, initially, may cause substantial damage to their hosts [16, 17]. The best-studied example of a horizontally transferred element is that of the *P-*element, a DNA transposon. *P-*element was acquired by *D. melanogaster* by horizontal transfer from a distant relative, *D. willistoni* [18], and spread rapidly through the species, as evidenced by its absence from old laboratory stocks collected before the *P-*element was widespread [19–21].

In its native species, and in current *D. melanogaster* populations, the *P-*element is not associated with any gross defects [18, 22]. But when males from strains collected after the invasion (‘P strains’) are crossed to females from old laboratory stocks (‘M strains’), the F1 offspring suffer a number of abnormalities, including recombination in males, increased mutation rates, frequent sterility, and abnormally small (dysgenic) gonads [16, 19, 23–25]. These abnormalities, collectively called ‘hybrid dysgenesis’, appear to be due to DNA damage caused by actively transposing *P-*element in the germline [17, 22, 26, 27]. In the reciprocal direction, the F1 offspring, though genetically identical, are normal. The reason is that these offspring are protected by piRNAs (<u>P</u>IWI <u>i</u>nteracting RNAs, a type of small RNA) homologous to the *P-*element, which are sequestered in the egg cytoplasm of P-type females. As these protective piRNAs target the *P-*element mRNA for degradation [28], their presence reduces *P-*element activity and the resulting DNA damage that lead to hybrid dysgenesis.

Since its initial discovery, *P-*element has been one of the most important tools in *Drosophila* genetics, providing the basis for genetic manipulations and mutagenesis (*e.g*., [12, 29]). The *P-*element had been thought to be absent from *D. melanogaster’s* close relatives [30], but recent work has detected it in *D. simulans*, apparently horizontally acquired from *D. melanogaster* [31]. As the *D. simulans* elements recovered are identical in sequence to a type that segregates at low frequency in *D. melanogaster*, and do not include the most common *D. melanogaster* element, this acquisition is likely due to a single transfer event [31]. Here, we report a hybrid dysgenesis-like phenotype in crosses between strains from different *D. simulans* populations, and show this phenotype occurs in crosses between strains collected before and after a rapid invasion of the *P-*element.

## Results

### Hybrid dysgenesis in *D. simulans*

We assayed for a hybrid dysgenesis-like phenotype in *D. simulans*, using the absence of developed ovaries as an indicator (Fig 1 and S1 Fig, see also [24, 32]). We considered a pair of strains to show hybrid dysgenesis when the F1 females from reciprocal crosses show a significant difference in the number of dysgenic ovaries [with significance defined as *p* ≤ 0.05 in a Fisher’s exact test (FET)]. Differences between F1 females from reciprocal crosses, which have identical nuclear genotypes, are likely due to different maternal contributions to the egg cytoplasm and implicate a cyto-nuclear incompatibility, as in the *D. melanogaster* hybrid dysgenesis system [19, 28].

**Fig 1.**
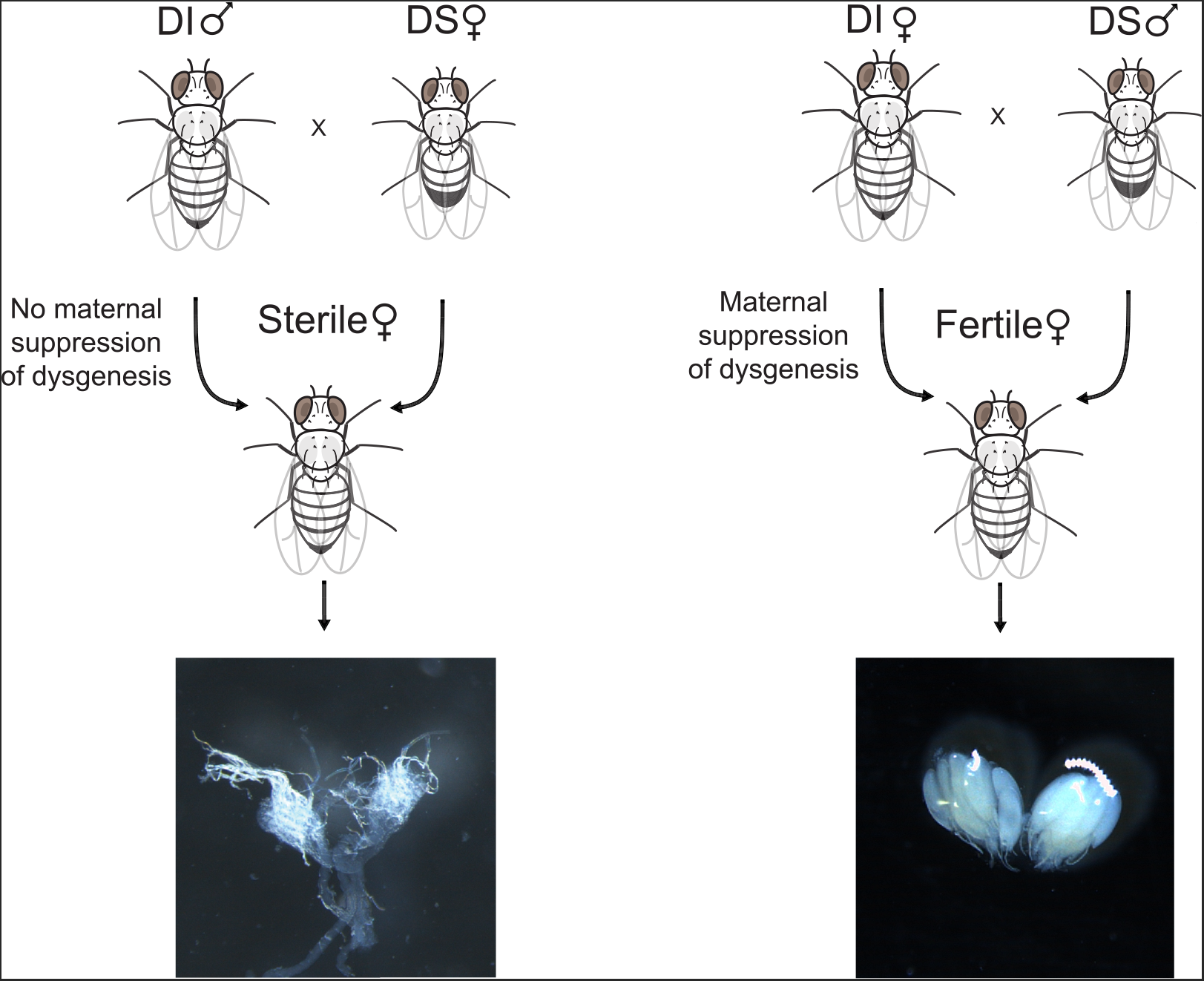
Hybrid dysgenesis in *D. simulans*. Outcome of a reciprocal cross between *D. simulans* strains. In the non-dysgenic direction of the cross (left, FL31♀ x M252♂), most female offspring have normal ovaries, while in the reciporical, non-dysgenic cross (right, FL31♂ x M252♀), many female offspring suffer from malformed ovaries.

For our initial survey, we used 22 strains collected in Georgia in 2009 (generously donated by P. Haddrill) and 10 strains collected in Madagascar in 2004 & 2008 (collected by B. Ballard and J. David; S1 Table). We paired strains from the two populations randomly, but ensured that each strain was involved in multiple crosses (at least 5 for the Madagascar strains and at least 2 for the Georgia strains). Among these crosses, 38.7% (36 of 93) showed unidirectional gonadal dysgenesis in F1 females at 29°C, a temperature which induces hybrid dysgenesis in *D. melanogaster*, but not at 25°C (S2 Fig). In addition to transposable elements, cyto-nuclear incompatibilities can also be caused by the intracellular bacteria *Wolbachia*. This is particularly likely here, as Georgia lines harbor a *Wolbachia* strain [33, 34] which appears to induce embryonic lethality in the offspring of crosses to Madagascar males. When we repeated the crosses after curing the strains of their *Wolbachia* infections, however, we found qualitatively similar results for infected and uninfected versions of the strains (S2 Fig, FET on data from the same cross performed with cured and uncured strains, *p* > 0.05 for all).

Inspection of the data shows that dysgenic crosses almost always involved one of five Georgia lines as the paternal line (S2 Fig). Control crosses between these “dysgenesis-inducing” (DI) lines showed no significant unidirectional dysgenesis, nor did crosses of these DI lines to 12 of the other Georgia lines (a subset of crosses are shown in S2 Fig C and D). In contrast, all 10 strains from Madagascar were “dysgenesis-susceptible” (DS) when crossed to DI lines; crosses between these lines also resulted in no hybrid dysgenesis (S2 Fig). The DI and DS classes are analogous to the P-type and M-type of *D. melanogaster*, respectively.

Using the Georgia and Madagascar strains with known DI and DS phenotypes, we unambiguously cytotyped 160 additional strains (cured of *Wolbachia*) from 23 populations sampled between 2002 and 2014 (Fig 2 and S3 Fig, S1 Table). We reciprocally crossed each strain to at least one known DI and DS strain, and roughly classified tested strains based on the results of the four crosses (see Materials and Methods). In addition to DI and DS types, we also observed a third type of strain, “dysgenesis-resistant” (DR), which produce no dysgenic offspring in either cross. These correspond to the Q types in *D. melanogaster*, which repress transposition of the *P-*element, but which contain few or no active elements [25, 35]. In fact, the resistant strains probably encompass a range of intermediate cytotypes (*i.e.*, corresponding to the M’, Q and weak P-types in *D. melanogaster* [17, 25, 35–38]). It is likely that these strains contain copies of the dysgenesis-causing element, but in degraded form, at low copy number, or with weak expression. These copies may nevertheless be able to produce piRNAs suppressing the TE [28, 39].

**Fig 2.**
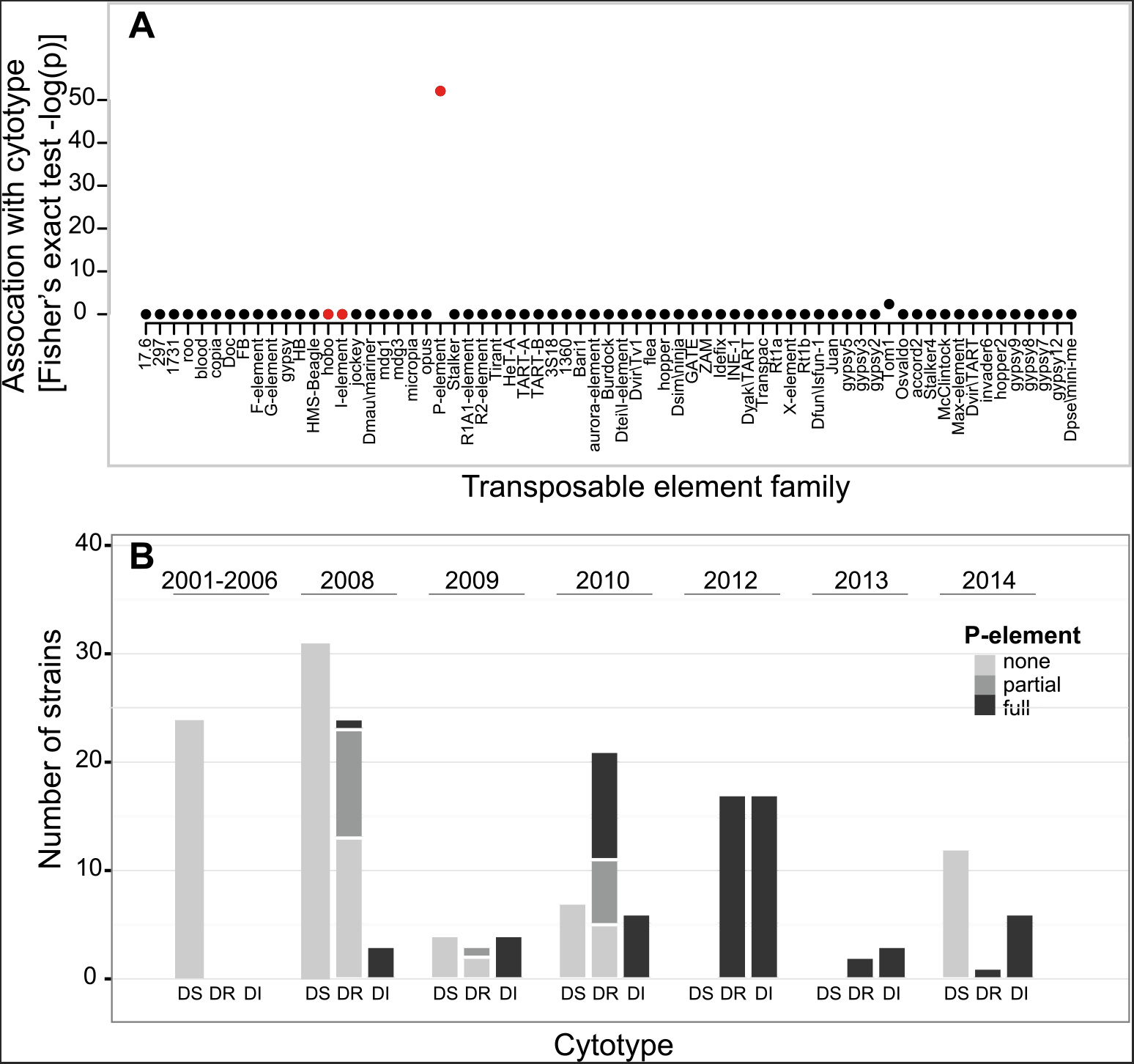
Cytotypes of lines assayed for dysgenesis. **A)** Plot showing the significance of association between the presence of different TE families in lines and their ability to cause hybrid dysgenesis (-log(p) from Fisher’s exact tests). TEs that previously have been associated with hybrid dysgenesis (*P*-element, *hobo*, *I*-element) are highlighted in red. **B)** The cytotypes and *P-*element infection status of a worldwide sample of *D. simulans* lines is shown. Populations from the same year are grouped together by cytotype. Two strains showed amplification of all four exons individually, but not of full-length element.

Overall, 19.3% of strains were DI, 26.5% DR, and 54.2% DS. Fertility assays performed on a subset of these crosses show that F1 females from dysgenic crosses show higher rates of sterility and produce fewer offspring than their genetically identical sisters from reciprocal crosses (S4 Fig; Mann-Whitney U test, *p* < 0.05).

### The *D. simulans* hybrid dysgenesis phenotype is associated with the P-element

Given the similarity between transposable element-induced dysgenesis in *D. melanogaster* and the results from the *D. simulans* crosses, we suspected that the DI lines might have a dysgenesis-inducing transposable element that is absent from the DS lines. Since the initial discovery of *P-*element, two other hybrid dysgenic systems have been discovered in *D. melanogaster*, the hobo-empty and inducer-reacter systems [40–42] and one in *D. virilis*, at least partly due to the Penelope element [43]. The existence of other dysgenesis-inducing TEs suggests that *P*-element might not be the cause of the hybrid dysgenesis seen here.

To identify TE families potentially associated with the dysgenesis-inducing agent, we used RNAseq data collected from pooled lines from a Florida population (originally collected in 2010, data from [31]), which shows both DI and DR cytotypes. We found 68 expressed TE families in these data. We investigated whether any of these elements appear to be associated with the observed hybrid dysgenesis, using three sources of evidence. First, we used genome sequence data from 12 single F1 females, produced by crossing males from the Florida population (4 DI strains, 6 DR strains & 3 DS strains collected in Florida 2010) to females from a single Madagascar strain, M252 (a DS strain, collected in Madagascar in 2004 [44], which lacks *P*-elements). We examined these sequences for TE families present in the Florida lines, but absent from M252. Only one of the expressed TE families showed an appreciable level of coverage and differed between the F1 flies and the M252 line— the *P-*element (S2 Table and S5 Fig). With short-read sequence data, we could not unambiguously identify full-length insertions, but at least three of the sequenced strains showed coverage of the complete *P-*element reference sequence [31].

Second, we looked for an effect of *P-*element copy number on the level of dysgenesis, as strains with more *P*-element copies are likely to suffer from more DNA damage in the presence of active transposase than those with fewer copies (see [16]). As a proxy for copy number in the Florida lines, we used sequence coverage of the complete P-element reference in the genome sequence data from the 12 F1 Florida X M252 females. Consistent with expectations, the strength of dysgenesis increases with coverage of the *P-*element in crosses where the male parent is from Florida [with coverage of the *P-*element standardized by the average coverage of the euchromatic genome; S6 Fig; binomial <u>g</u>eneralized <u>l</u>inear <u>m</u>odel (GLM); *z* = 6.49, *p* = 8.49 x 10^-11^]. There is no relationship between dysgenesis and coverage for the reciprocal cross (*z* = -1.128, *p* = 0.26), or for coverage of other TEs in any cross (*p* > 0.05 for all, S2 Table).

Finally, we designed primers within the exons of the 68 expressed TEs (S2 Table), and tested for differences between previously cytotyped DI and DS strains using PCR. All but two of the elements with appreciable coverage were found to be either present in or absent from all strains, regardless of cytotype (*P-*element and *Tom1*; S2 Table). *Tom1* is not significantly associated with cytotype (present in 35 of 35 DI strains, and 71 of 78 DS strains, FET *p* = 0.097), and there was no relationship between sequence coverage of *Tom1* and the strength of dysgenesis, as described above (S6 Fig; binomial GLM; *z* = -0.551, *p* = 0.581). In contrast, the presence of *P-*elements is strongly associated with the DI cytotype, with products corresponding to *P-*element (a subset of which were verified by Sanger sequencing) amplified from all 35 DI strains. We were also able to amplify full-length *P-*element from 31 of these strains using standard PCR, and from the 4 remaining strains using RT-PCR — we were able to amplify all 4 exons of the *P-*element individually in all of the 35 strains. In contrast, no *P-*element could be amplified from any of the 98 DS strains (full-length element obtained using standard PCR in DI vs. DS strains; FET *p* < 2.2 x 10^-16^). Using RT-PCR, we confirmed that full-length *P-*element is expressed in the 5 DI strains used in experimental crosses for cytotyping, and that no full-length transcript could be amplified from the DS strains used (S7 Fig A).

Based on these three lines of evidence, we conclude that the *P-*element is the likely cause of the dysgenesis we see in *D. simulans*. In fact, the *P-*element is known to be active and cause dysgenesis in *D. simulans* when artificially introduced [32, 45]. Consistent with a homology-based silencing mechanism, such as that mediated by piRNAs [28], maternal suppression of dysgenesis appears to be associated with the presence of the *P-*element in some form, either partial or full-length: we were able to amplify *P-*element sequence from all 48 DR strains (48 DR strains with *P-*element *vs*. 98 empty DS strains; FET *p* < 2.2 x 10^-16^). In 18 of the 48 DR strains, we found only partial copies of the *P*-element, consistent with the production of piRNAs from degraded copies of the transposon [46, 47]. If, as in *D. melanogaster*, the maternal suppression of *P-*element sequence in a bi-directional piRNA cluster (one to which both sense and antisense piRNAs map) [46–48], the resistant strains might express both sense and anti-sense transcripts). We performed RT-PCR for each direction of the transcript separately on 10 strains (3 DI, 4 DR, 3 DS), and, as expected, found expression for both the sense and anti-sense RNA strands for DI and DR lines, but for neither direction in the DS lines (S7 Fig B).

### The *P*-element invasion of *D. simulans* occurred rapidly on three continents

We reasoned that, as in *D. melanogaster*, the DS and DI strains might show temporal and/or geographical differences [20, 21, 49–51]. In fact, the frequency of the DI and DR dysgenesis types increase over time (binomial GLM; Africa: *z* = 5.822, *p* = 5.81x 10^-9^), and previous work shows differences in *P-*element content between a samples from 2010 and 2012 [31]. We tested a collection of 631 samples (isofemale lines, ethanol-preserved flies, and DNA extracted from wild flies) collected from 54 locations between 1984 and 2014 for the presence or absence of the *P-*element (S1 Table, Figs 3 and 4). To detect *P*-element, we used PCR, as before, with multiple sets of primers and repeated independent DNA extractions (see Materials and Methods). As before, the primers were used to amplify products that correspond to the full-length element and to each of the four exons. As for the cytotyped lines, we found strains with full-length copies, strains with only partial copies (in which at least one exon consistently failed to amplify), and strains with no successful amplification of any of the tested *P-*element fragments. The partial copies were usually missing internal exons, a common by-product of imperfect excision events in *D. melanogaster* [16, 52]. Strains with full-length elements often also contained other, partial copies with similar internal deletions.

**Fig 3.**
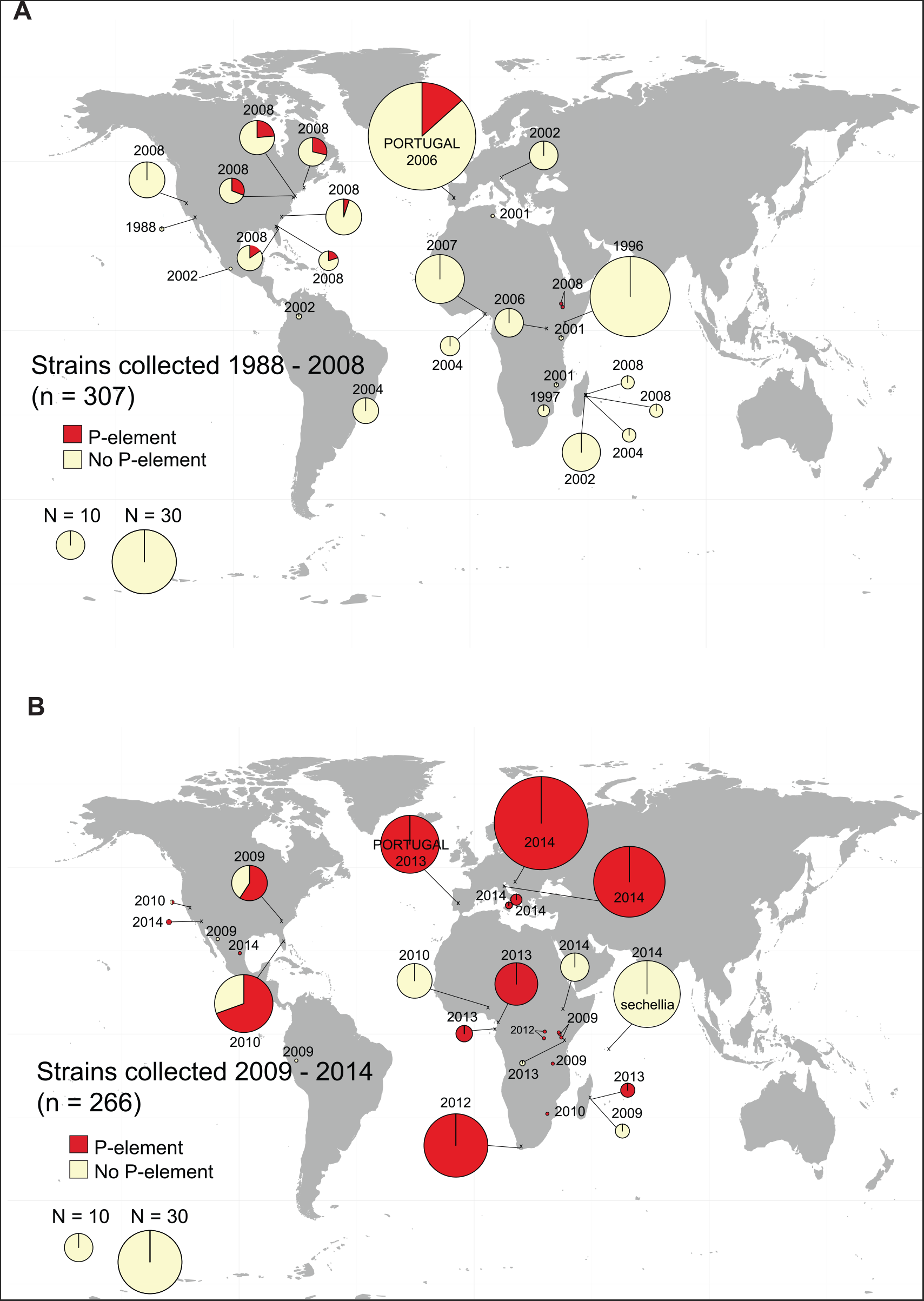
Partial and full *P-*element copies in *D. simulans* collected A) 1984-2008, and B) 2009-2014. Map shows the approximate location where strains were collected, and pie charts show the proportion of strains containing *P-*element (either full-length copies or individual exons could be amplified by PCR) and no copies (no *P-*element of any kind could be sampled). The area of the pie charts is proportional to the number of strains sampled (see Legend), except for single points, which indicate one strain.

**Fig 4.**
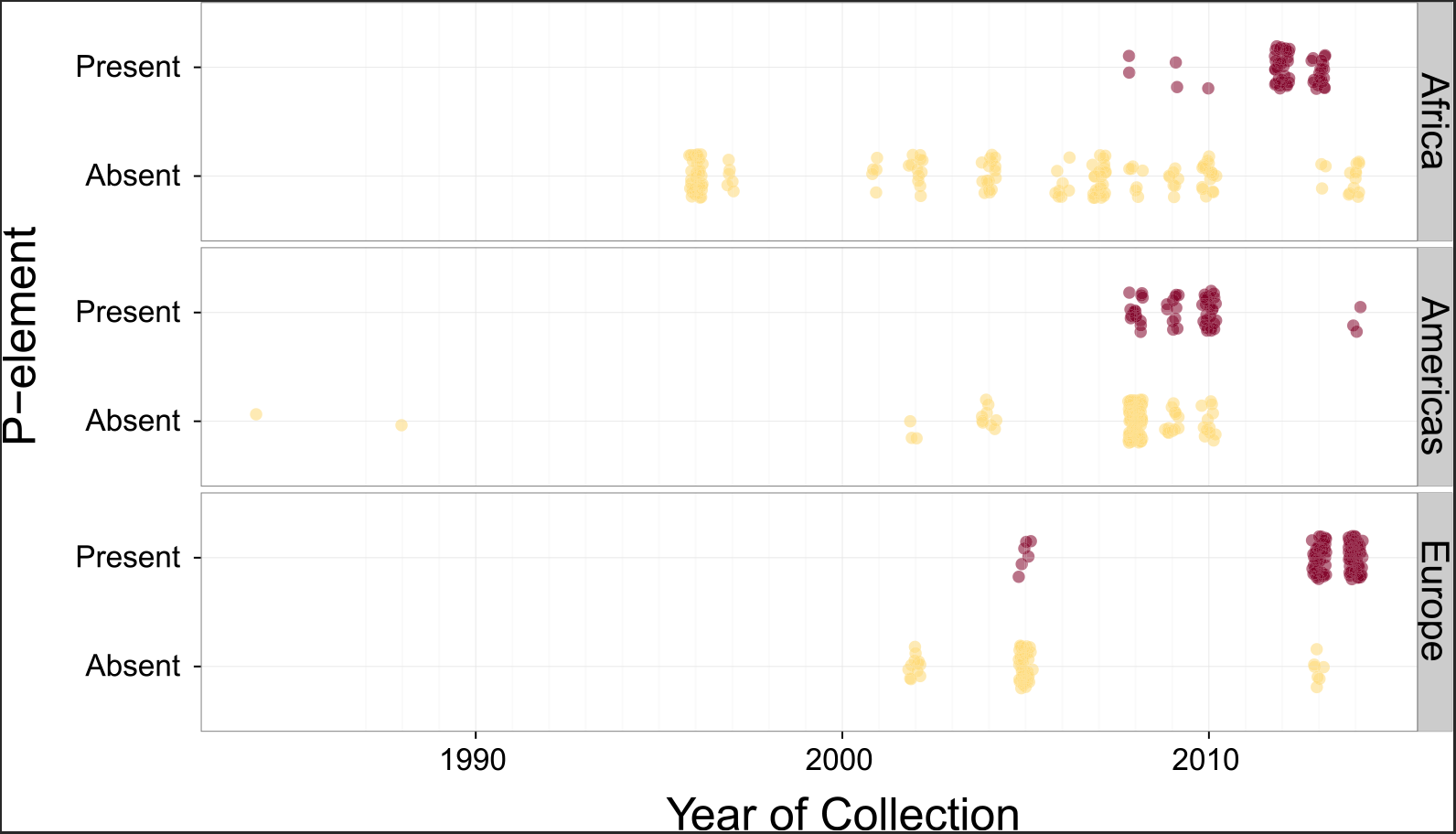
Time series of *P-*element in different forms in *D. simulans*. The plot shows strains with full (and partial) and no *P-*elements over time in Africa (top panel), North and South America (middle panel), and Europe (bottom panel) across time. Each point indicates a strain, with overlapping points jittered slightly. Data from the 1980’s in the Americas are from Brookfield (1984) [30].

The fraction of strains containing the *P-*element increases over time on all continents surveyed. In collections from 2004 to 2009, strains free of the *P-*element predominate (Fig 3A). After 2009, most strains harbour some form of *P-*element, and many have full-length elements (Fig 3B). *D. sechellia*, a close relative of *D. simulans* with which it is known to hybridize [53, 54], remains *P-*element free in our 2014 samples (kindly provided by D. Matute, Fig 3B). We find a significant association between when a strain was collected and whether it contains any *P-*element copies, regardless of the geographic region from which it was collected (Fig 4, binomial GLM with year, geographical region, and an interaction term fit to the *P-*element for each strain; year as *z* = 5.218 *p* = 1.81x 10^-7^).

By 2013, almost all sampled strains contained *P-*elements in some form (Fig 3). It is formally possible that this apparent spread is instead due to the loss of *P-*element in strains propagated for several years in the laboratory. This process could not have occurred, however, in the Portuguese samples, for which genomic DNA was extracted from directly wild flies, and these show a similar pattern (compare Portuguese samples from 2006 and 2013 in Fig 3). The rapidity of the spread makes it difficult to identify a likely origin point for the invasion, as the *P-*element appears to have invaded all three regions nearly simultaneously (Fig 4), with statistical differences between regions (S3 Table) apparently due to low prevalence of the *P-*element in a few late samples, rather than pointing to a geographic origin of the invasion.

## Discussion

Here, we identified a hybrid dysgenesis-like system in *D. simulans*. Using next generation sequencing information and PCR, we find that the *P-*element, which has recently invaded *D. simulans* [31], is associated with the hybrid dysgenesis phenotype, and spread rapidly across Europe, Africa, and North America. In fact, compared to *D. melanogaster*, the spread of the *P-*element in *D. simulans* is surprisingly fast: In *D. melanogaster*, the first *P-*type fly was collected from the wild in 1954, and most wild strains were infected by 1974 [20]. We find the first evidence of the *P-*element in *D. simulans* samples from 2006, with *P-*elements appearing in nearly all lines sampled since 2013, seven years later (apart from one sample from Ethiopia; Fig 3, Fig 4, S1 Table).

There are a number of reasons the *P-*element invasion may have occurred more rapidly in *D. simulans* than in *D. melanogaster*. For one, *D. simulans* appears to enjoy lower rates of *P-*element induced gonadal dysgenesis than *D. melanogaster;* high rates of sterility are expected to hamper the spread of the *P*-element by reducing the fitness of its carriers. Direct comparisons between the species are problematic, as the severity of hybrid dysgenesis increases with copy number, which differs between the species ([19, 24, 26, 31]; S2, S4, S6 Figs). But in comparisons between artificially constructed *D. melanogaster* and *D. simulans* strains with comparable copy numbers, P-element does appear to be less active in *D. simulans* [32, 45, 55].

Another, simpler explanation for the difference in the speed of invasion is higher migration rates between *D. simulans* populations than for *D. melanogaster* as they experienced the *P-*element invasion. Such a scenario seems reasonable: Unlike *D. melanogaster, D. simulans* cannot over-winter in temperate regions, and instead reinvades the northern limits of its range annually. Sequence data correspondingly suggests a high rate of homogenization between *D. simulans* populations at nuclear loci [56–61]. Furthermore, for both species, migration might be largely human-aided, and the global movement of fruit has increased dramatically in the years since the spread of the *P-*element in *D. melanogaster*. That is, global imports and exports of fleshy fruits (measured in tons) grew about 4% per year from 1961-2013 (the years tracked by the Food and Agriculture Organization of the United Nations Statistics Division). As a result, the average tonnage of fruit imports was 5-fold higher in the years measured during the spread of the *P-*element in *D. simulans* compared to *D. melanogaster* (for imports, 1961-1974: 12.5 vs. 2006-2014: 61.5 million tons/year; exports tracked imports closely). While these statistics capture only the movement of fruit between countries, the increase in movement seems likely to have boosted the global spread of the *P-*element in later years.

The *P-*element in *D. simulans* adds to a handful of documented cases of the spread of a horizontally transmitted element through a *Drosophila* species [33, 62–65]. Several of the other cases include intracellular bacteria, which are capable of behaving selfishly, but in these cases also appear to be conferring a benefit to their hosts [64–67]. In contrast, the *P-*element appears to be spreading mainly *via* a selfish mechanism; though the occasional insertion may be beneficial [22, 68]. In response, both *D. simulans* and *D. melanogaster* have quickly evolved to suppress the dysgenesis phenotype by the end of the invasion (present study, [20, 38]), suggesting strong selection to ameliorate the the P-element’s worst effects, and underscoring the evolutionary challenge posed by selfish elements.

## Materials and Methods

### Fly strains

We used *D. simulans* isofemale lines collected across multiple locations over 15 years (see S1 Table for details). Flies were reared on molasses–yeast–agar *Drosophila* medium at 18°C unless otherwise stated. The species identity of all lines was confirmed either by visual inspection of male genitalia or, when this was not possible, by amplification of PCR products that yield species-specific product lengths (using primer sequences for Argonaute2, FBgn0087035, kindly provided by D. Obbard: Mel_Sim_F 5’-CCCTAAACCGCAAGGATGGAG-3’, D_sim_303_R 5’- GTCCACTTGTGCGCCACATT-3’,D_mel_394_R5’CCTTGCCTTGGGATATTATTAGGTT-3 ‘).

Unless otherwise stated, all lines were cured of *Wolbachia* infections before crosses were performed, in order to obtain sufficient offspring from crosses that normally show high levels of cytoplasmic incompatibility. (If known, the infection status of lines prior to curing is given in S1 Table.) To cure lines, we raised larvae on food supplemented with 0.05mg/ml tetracycline-hydrochloride for two generations. After curing, flies were tested for *Wolbachia* using PCR, with primer sequences wsp81F (5’-TGGTCCAATAAGTGATGAAGAAAC-3’) and wsp691R (5’-AAAAATTAAACGCTACTCCA-3’), and a cycling program of 94°C for 5’ followed by 30 cycles of 94°C 30”, 55°C for 60”, 72°C for 2’, and ending with one cycle of 72°C for 10’, as in [69].

*Hybrid dysgenesis assays*. To assay for hybrid dysgenesis, we performed reciprocal crosses between lines using 5 virgin males and females from each line. Flies were left to lay eggs for a total of 9 days at 29°C, transferred to a fresh vial every 3 days, and offspring reared at 29°C (28°C is the lowest temperature at which dysgenesis is observed in *D. melanogaster* [19, 24]). We dissected 30 F1 females from each cross, and assessed them for the presence or absence of two normal ovaries. We scored the number of females lacking one or both ovaries as dysgenic, following [24], and considered crosses showing a significant difference (at the 5% level) to be dysgenic.

To classify strains as DI, DR, or DS, we reciprocally crossed each strain to at least one known DI and DS ‘tester’ strains, and roughly classified tested strains based on the results of the four crosses. Specifically, we defined strain as:

> DI, if strain x tester DI, non-dysgenic; strain x tester DS, dysgenic
>
> DS, if strain x tester DI, dysgenic; strain x tester DS, non-dysgenic
>
> DR, if strain x tester DI, non-dysgenic; strain x tester DS, non-dysgenic.

Some strains could not be unambiguously classified (5 borderline DI/DR types, and 6 borderline DR/DS types, and 3 that tested as both DI and DS). These strains were not used in the analysis; in all, 181 strains were unambiguously cytotyped.

*Sterility Assays*. To perform the sterility assays shown in S4 Fig, we reciprocally crossed lines as described previously at 29°C. For each line, individual virgin F1 females from both reciprocal crosses were mated to control virgin males (raised at 25°C) at 25°C. Flies were left to mate and lay eggs for a total of 7 days and then removed. The offspring from these crosses were collected and counted every 2 days. As fertility of F1 females from PxM crosses in *D. melanogaster* recovers with age [70], we checked the fertility of young (3-9 days old) and old (10-16 days) females separately. We compared the fertility of females from the ‘dysgenic’ and ‘non-dysgenic’ direction of the crosses using a Mann-Whitney U test.

### Analysis of sequence data

To find TEs expressed in the Florida lines, we used RNA-seq data form a pooled sample of lines from this population (accession number PRJEB7936 [31]). Paired end Illumina reads were mapped to a *D. simulans* reference genome produced from a Madagascar strain [44] and 179 *Drosophila* TE sequences (http://flybase.org/; [71, 72]) using GSNAP ([73]; parameters: -n =1, -N = 1).

We examined whole genome sequence from the Florida strains using pairedend Illumina reads (SRA PRJEB7936 [31], and PRJNA308281) collected from 12 barcoded individual F1 offspring of the M252 reference strain females (sequence data under SRX504933) crossed to males from Florida (2010) isofemale lines, and the paired end reads from the M252 reference strain [44]. We mapped these data to the *D. simulans* M252 reference genome using BWA-SW with the default parameters [74, 75] and following the mapping protocol described in [76]. We counted the number of insertions per individual sequenced in two ways: First, we used PoPoolationTE [74], with the requirement that each insertion be supported by at least 3 reads. Second, due to the fragmented nature of the *D. simulans* genome sequence, some insertion sites may not be recovered by PoPoolationTE, which relies on mapping reads to a reference. We therefore also estimated TE copy number per family by comparing average coverage of the TE sequence to average coverage of chromosome arm 2L.

### PCR assays

Primers for TEs were designed based on the EMBL sequences (S4 Table, http://flybase.org/; [71, 72]). The following PCR program was used: 94°C for 5’ followed by 30 cycles of 94°C 30”, 55°C for 60”, 72°C for 1-3’ (depending on the expected length of the PCR product, 1 minute per kilobase); 72°C for 10’. PCR products were run on a 1% agarose gel to test for the presence or absence of a TE. To survey for the *P-*element, primers were designed for each exon of *P-*element (S4 Table).

The presence of full *P-*elements was assessed using the forward primer for exon 0 and the reverse primer for exon 3, and using a primer annealing to both ends of the *P-*element (in the TIR region; sequence kindly provided by L. Teysset). To ensure that PCR failures are due to the absence of *P-*element, DNA quality was checked with a positive control PCR for an essential gene, and *D. melanogaster* Harwich P was used as a positive control in all PCRs. Sanger sequencing (performed by LGC Genomics, http://www.lgcgroup.com/) of a subset of PCR products was used to confirm that the amplicons correspond to *P-*element sequence (Genbank accession numbers KU719478-KU719502); the sequence of additional sequencing primers is available upon request. We required a minimum of two independent DNA extractions to consistent results for each line.

To perform RT-PCR, RNA was isolated from Florida, Georgia, Harvard, Maryland and Madagascar lines using Peqlab TriFast RNA isolation protocol, and extractions checked on a 1% agarose gel following treatment with DNase. We then performed RT-PCR using QIAGEN OneStep RT-PCR Kit, using the following program to detect full-length *P-*element RNA (using the forward primer for exon 0 and the reverse primer for exon 3): 45°C for 30’ for reverse transcription, then 95C for 15’, followed by 30 cycles of 94C for 15”, 55C for 60” and 68C for 6’; 68C for 10’ to amplify cDNA. Using the same program, reverse transcription was also performed separately for both forward and reverse of the *P-*element (using either the forward primer for exon 0 or reverse for exon 3) to generate unidirectional cDNA for PCR.

### Statistical analyses

All statistical analyses were performed using R [77]; https://www.rproject.org/). To estimate global trade in fruits, we downloaded data for total worldwide imports and exports of fruits and vegetables for the years 1961 to 2013 (measured in tons) from the FAOstat website (http://faostat3.fao.org/download/T/TP/E). From these, data for nuts, vegetables, and processed foods were excluded, resulting in data for 29 categories of fleshy fruit and berries. Using the *nls* function in R, we fit exponential growth models to these data of the form *y* = *a* exp(*bx*), where *y* is the exports in tons, *x* is the number of years since1961, and *a* and *b* are the estimated intercept and growth rate, respectively.

## Acknowledgements

Thanks to Viola Nolte for assistance with PCR, performing the sequencing and for providing information for many of the isofemale lines. We are grateful for helpful discussion provided by R. Kofler, W. Miller, K. Senti, and C. Vogl, and to B. Charlesworth, G. Lee, and three anonymous reviewers whose comments greatly improved the manuscript. We are especially grateful to all the researchers who generously shared fly collections (S1 Table), as this work would not have been possible without them.

## Supporting Information Legends

**S1 Fig.**
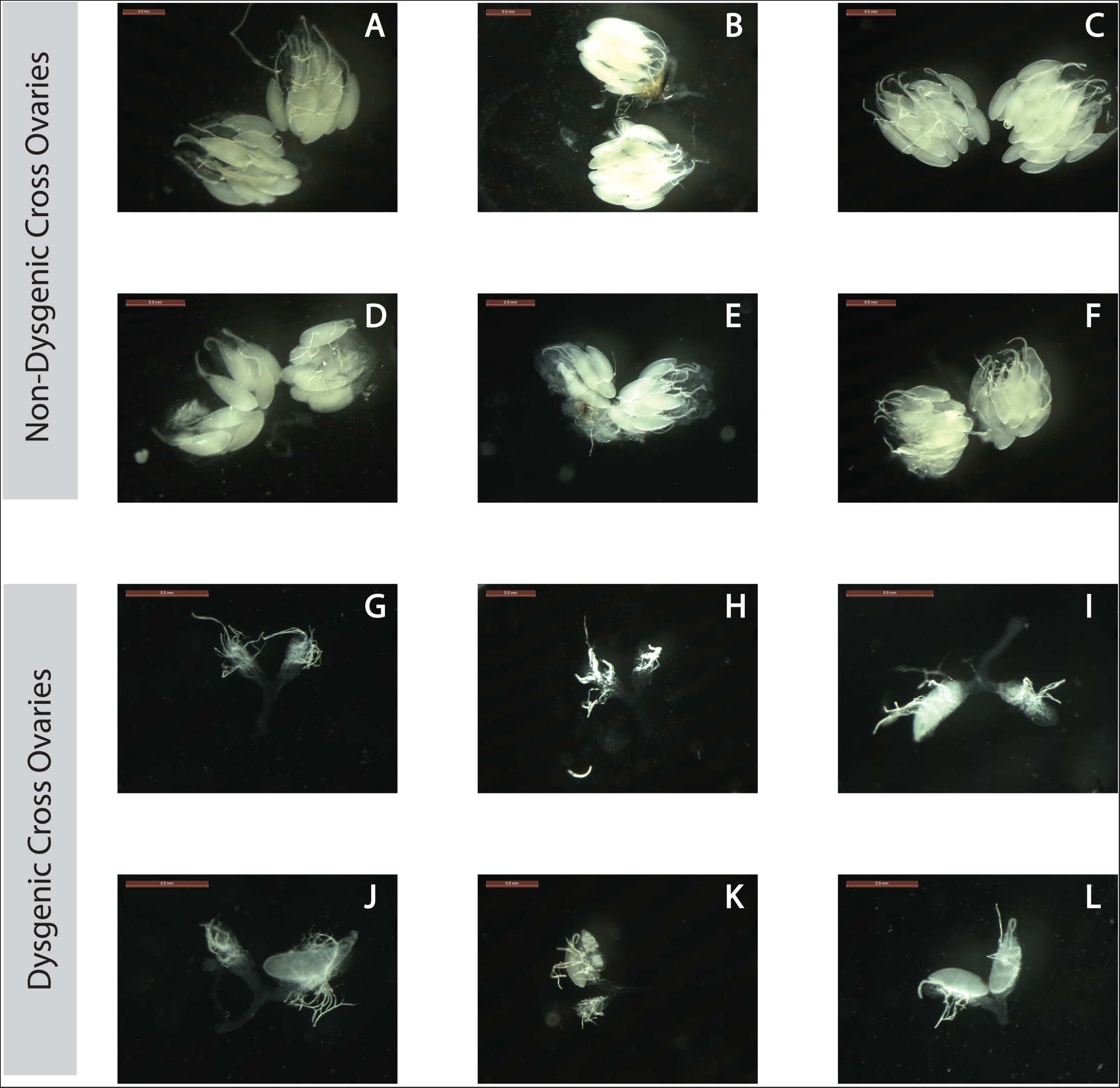
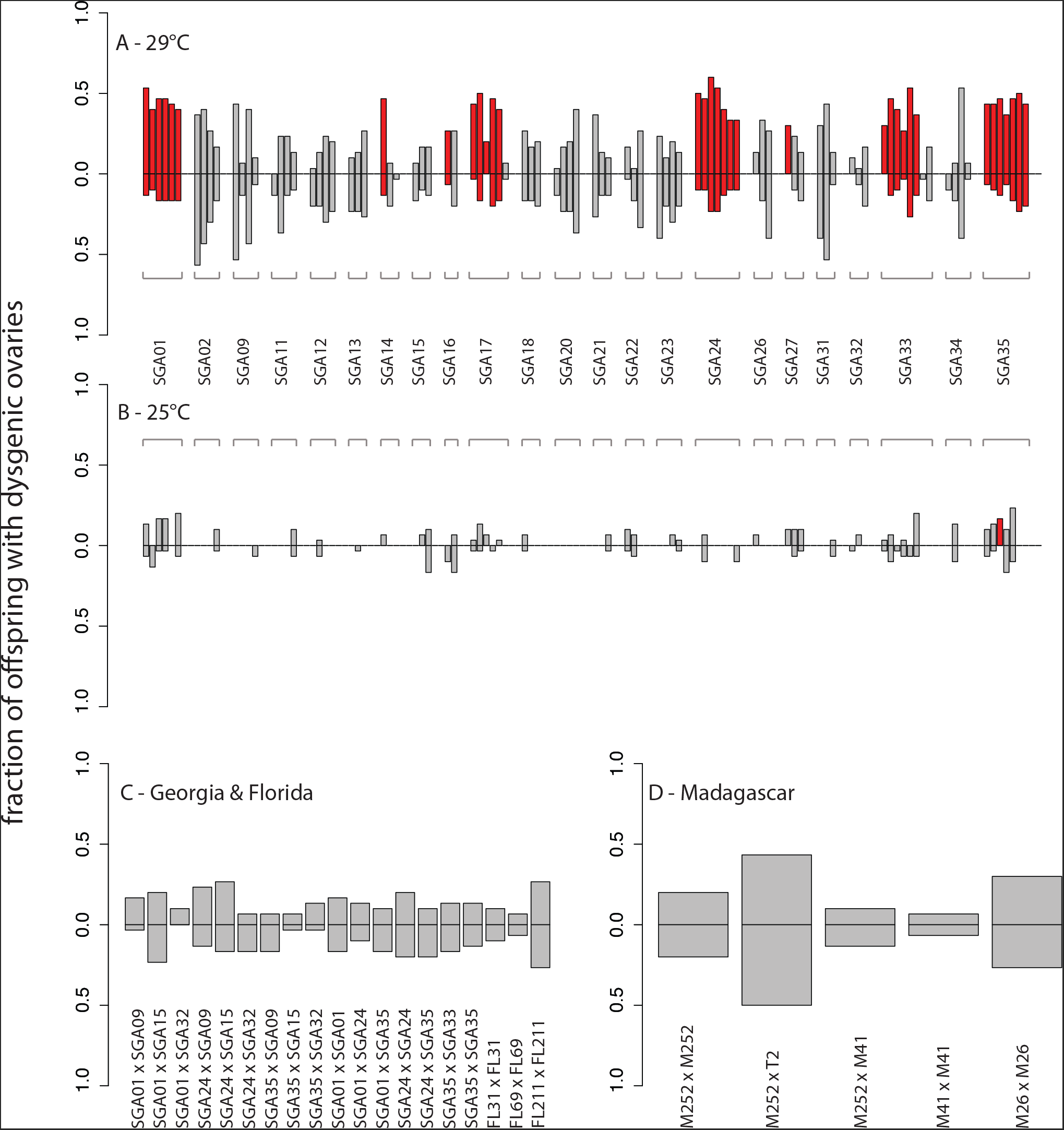
Dysgenic and non-dysgenic ovary dissections. Images showing non-dysgenic **(A-F)** and dysgenic ovaries **(G-L)**. The brown bar on each image is a scale of 0.5mm. All ovaries were taken from F1 females from crosses between SGA35 & M252 for each direction. Non-dysgenic ovaries, though they appear to vary in size and ovariole number (e.g. C vs. D), are much larger than dysgenic ovaries. In dysgenic ovaries, if present, they are usually rudimentary and only one or two ovarioles form properly (I-L). Dysgenic ovaries can also be asymmetric, as ovarioles will occasionally form on one side but not the other (I-K), as previously described in *D. melanogaster* [25]. Frequencies of total dysgenesis (G & H) vs. asymmetrical dysgenesis (I-L) vs. rudimentary ovaries (L) were not recorded during dissections, though total dysgenesis was seen in an estimated 70% of dysgenesis cases.

**S2 Fig.**
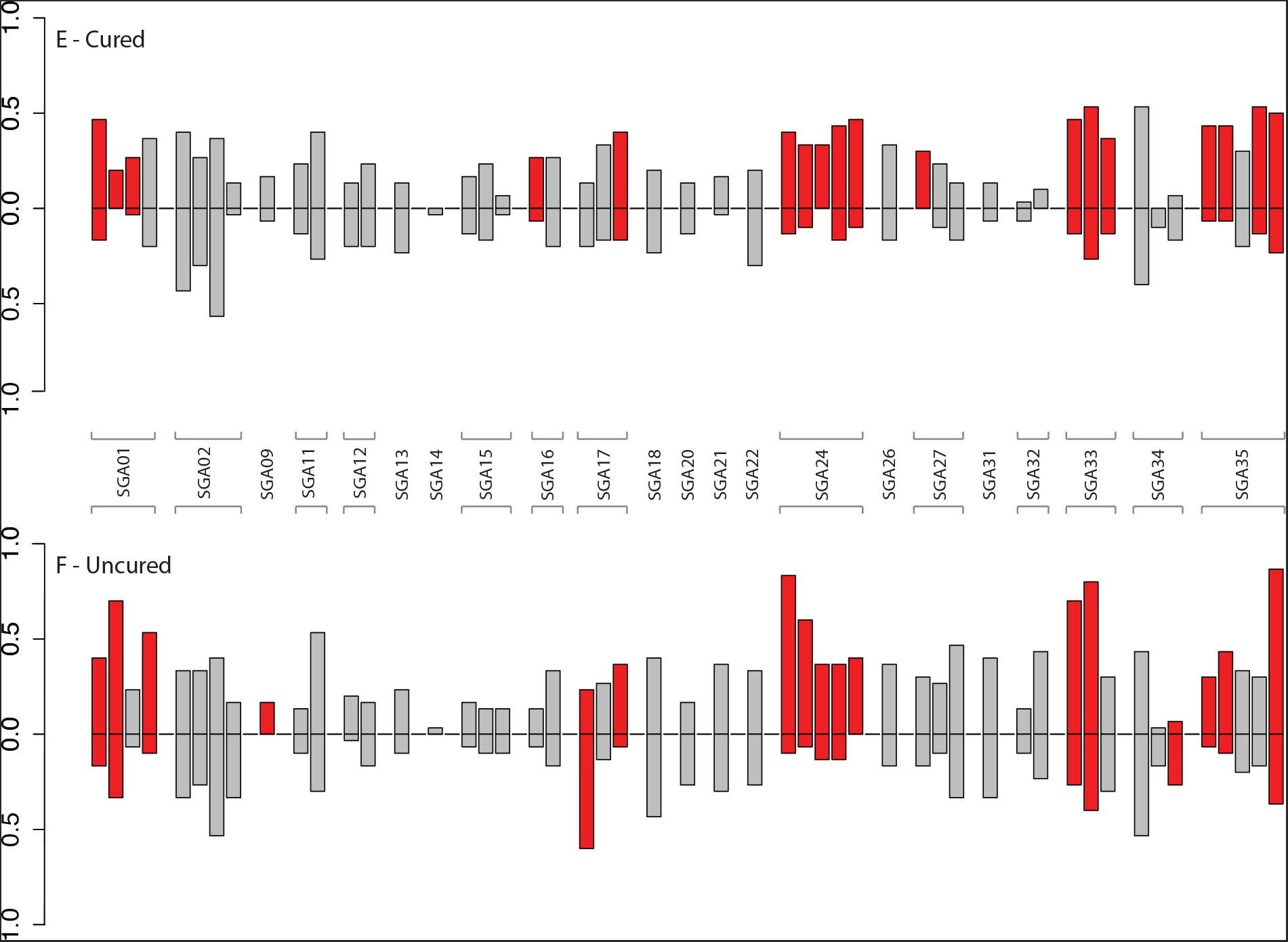
Hybrid dysgenic crosses and controls. Bar plots show the fraction of dysgenic offspring for both directions of each set of reciprocal crosses. Bars are coloured red when a significant difference between each reciprocal cross is found (Fisher’s Exact Test, *p* < 0.05). **A)** Initial crosses between Georgia (2009) and Madagascar (2004) flies at 29° C, with the fraction of dysgenic F1 females shown when the Georgia strain is paternal (positive direction) or maternal (negative direction). **B)** A subset of the crosses in panel A were repeated at 25° C, a non-dysgenic temperature in *D. melanogaster;* repeated crosses are shown in the same position as in panel A. **C)** Fraction of dysgenic offspring from crosses within and between Georgia and Florida lines at 29° C. The first strain named in the cross is the paternal strain in the positive direction and the maternal strain in the negative direction. **D)** Fraction of dysgenic offspring from crosses within and between Madagascar lines at 29° C. The first strain named in the cross is the paternal strain in the positive direction and the maternal strain in the negative direction. **E)** Fraction of dysgenic offspring from crosses between Georgia and Madagascar lines at 29C after curing of *Wolbachia* with tetracycline hydrachloride for 2 generations. **F)** Fraction of dysgenic offspring from crosses between Georgia and Madagascar lines at 29C before curing.

**S3 Fig.**
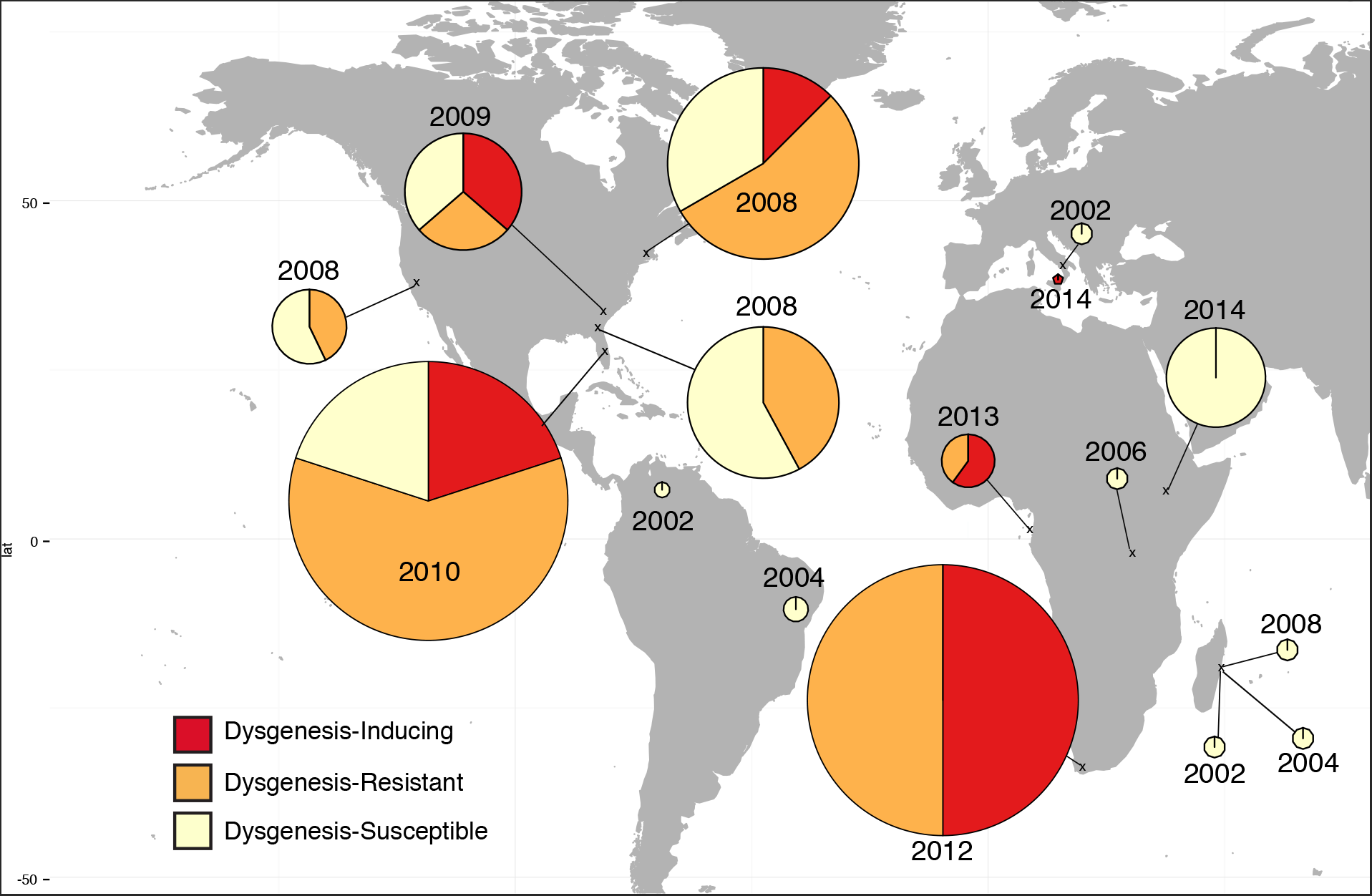
Hybrid dysgenesis map. Map shows the approximate location where strains were collected; pie charts show the proportion of each population which were dysgenesis-inducing (red), dysgenesis-resistant (orange) and dysgenesis-susceptible (yellow). This summarizes the data given in Table 2. The area of the pie chart is proportional to the number of strains sampled.

**S4 Fig.**
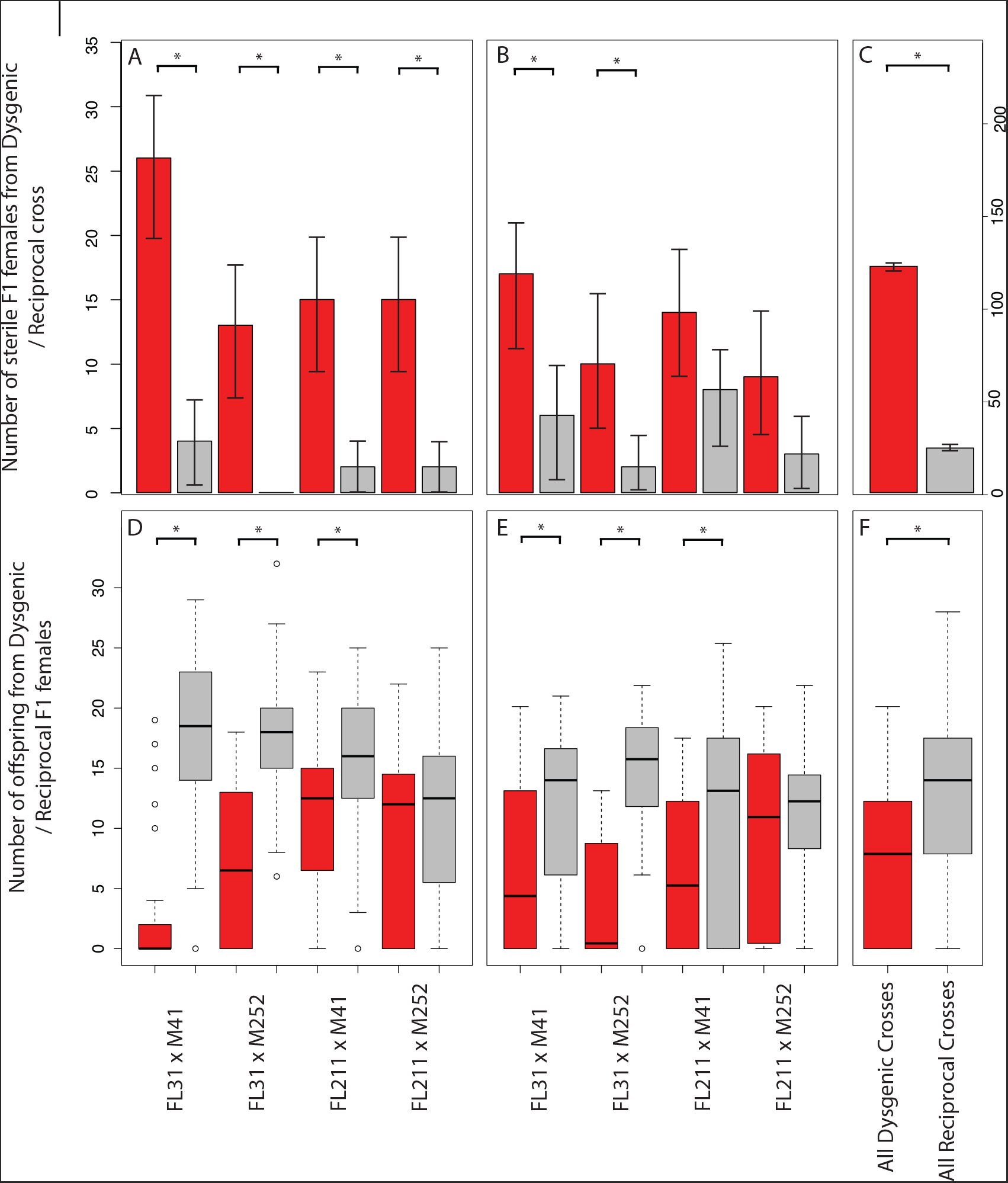
Sterility assay. Plots **A-C** show the number of sterile F1 offspring produced from the dysgenic (red) and reciprocal (grey) crosses (from 35 total offspring), shown with 95% confidence interval. Crosses that show a significant difference in sterility between directions are marked with a star (Fisher’s Exact Test *p* < 0.05). Plots **D-F** show the number of offspring produced by the female offspring from both the dysgenic and reciprocal cross. Crosses that show a significant difference in the number of offspring for each direction are marked with a star (Wilcox Rank Sum Test *p* < 0.05). Female parents were dissected after crossing to confirm the presence of gonadal dysgenesis (mean of 50.6% of female parents were dysgenic, compared to a mean of 8.6% dysgenic female parents in the reciprocal cross, note this is a similar proportion to the proportion of sterile F1 females). **A & D.** Females were mated after 3-9 days of aging. **B & E.** Females were mated after 10-16 days of aging. **C & F.** Combined results for all crosses and ages.

**S5 Fig.**
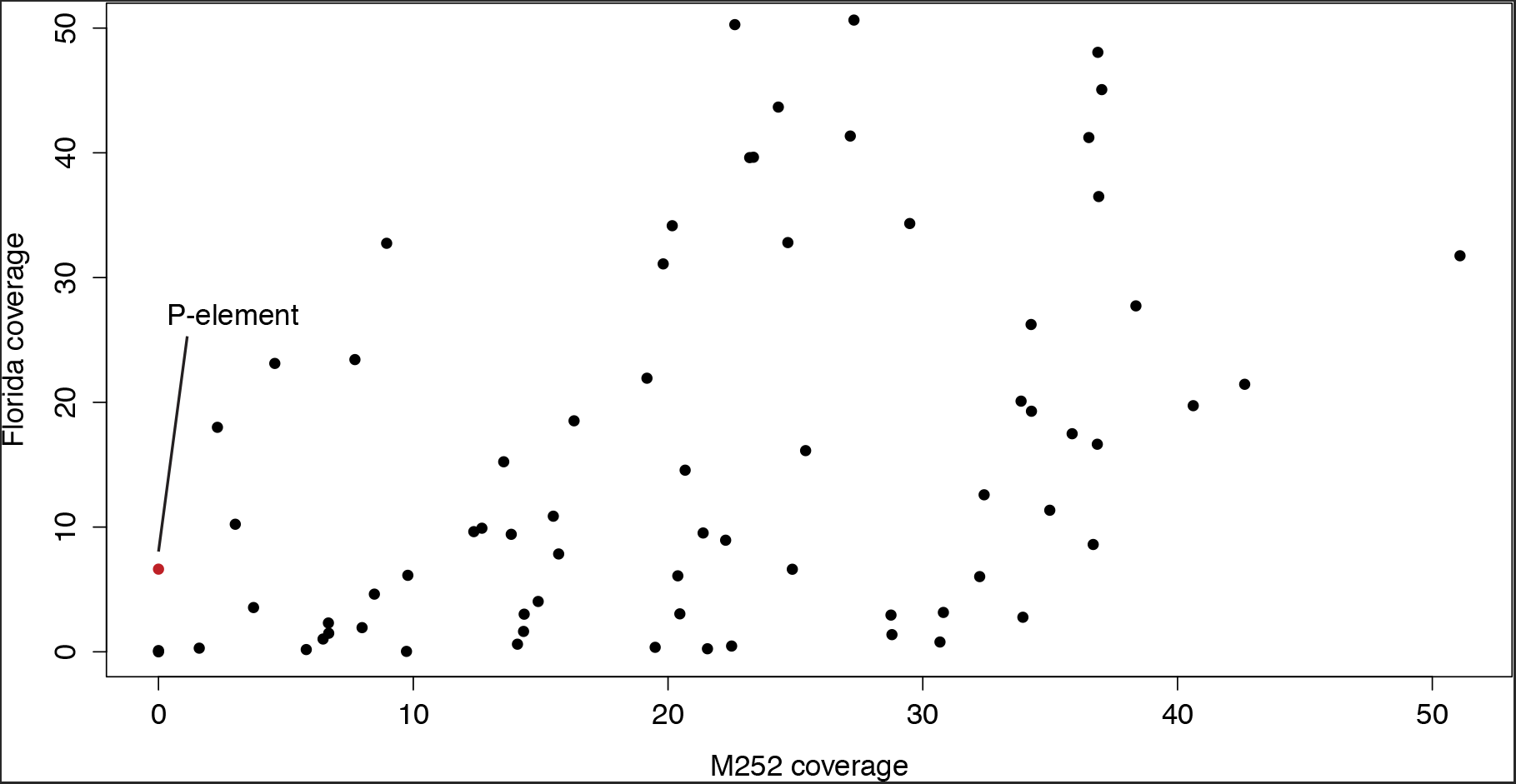
TE coverage comparison. Average coverage for TEs in the sequenced Florida (2010) x Madagascar (2004) isofemale lines *vs*. coverage for TEs in M252 (2004). Here, coverage is defined as the average per base coverage of TE sequence, divided by the average per base coverage of chromosome 2R in the line. The Florida coverage is taken as the average coverage of 12 available lines (FL3, FL6, FL98, FL101, FL116, FL136, FL168, FL174, FL189, FL198, FL208 & FL211). The Florida data (SRA:PRJEB7936, PRJNA308281) and Madagascar data (SRA:SRX504933) was mapped to the *D. simulans* reference genome alongside Flybase TE sequences to identify differences in coverage for these TEs. Only one TE was found in Florida but not in Madagascar, P-element.

**S6 Fig.**
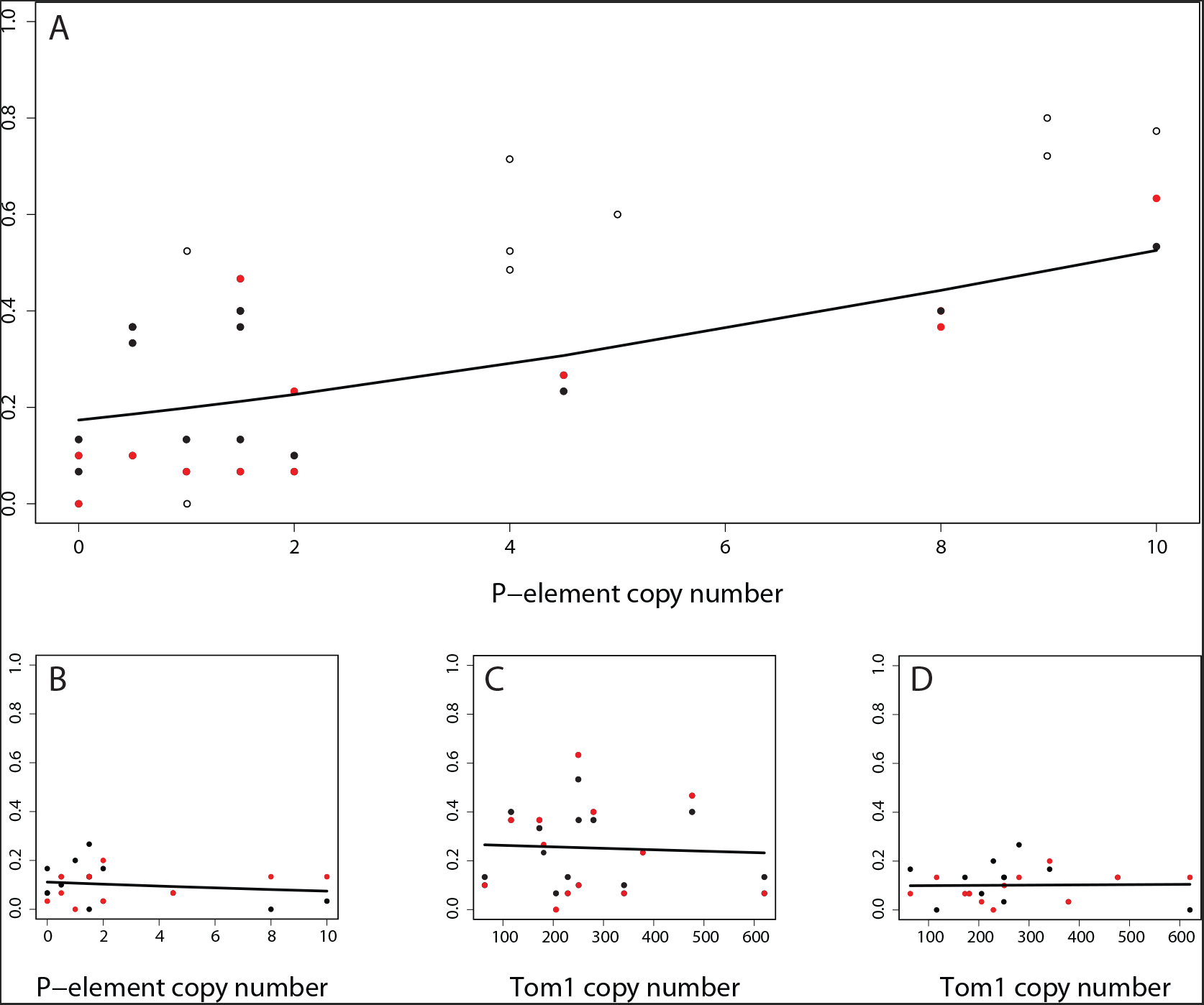
TE copy number vs. strength of hybrid dysgenesis. Estimated copy number of TE (estimated via dividing the average coverage of the TE in a sample by the average coverage of chromosome 2R) *vs*. the fraction of dysgenic offspring observed in a cross (12 lines from Florida population [described in S5 Fig] crossed to M26 [in red] and M252 [in black]), with lines showing the fit of a binomial generalised linear model. **A)** The correlation between the copy number of P-element and the proportion of dysgenic offspring from the paternal Florida crosses (S2 Table; z value = 6.49, *p* =8.49e-11). The equivalent association between dysgenesis and P-element copy number for *D. melanogaster* is shown with clear points [16]. **B)** The correlation between the copy number of P-element and the proportion of dysgenic offspring from the maternal Florida crosses (S2 Table; z value = -1.128, *p* = 0.259). **C.** The correlation between the copy number of Tom1 and the proportion of dysgenic offspring from the paternal Florida crosses (S2 Table; z value = -0.551, *p* =0.581). **D.** The correlation between the copy number of Tom1 and the proportion of dysgenic offspring from the maternal Florida crosses (S2 Table; z value = 0.135, *p* =0.893).

**S7 Fig.**
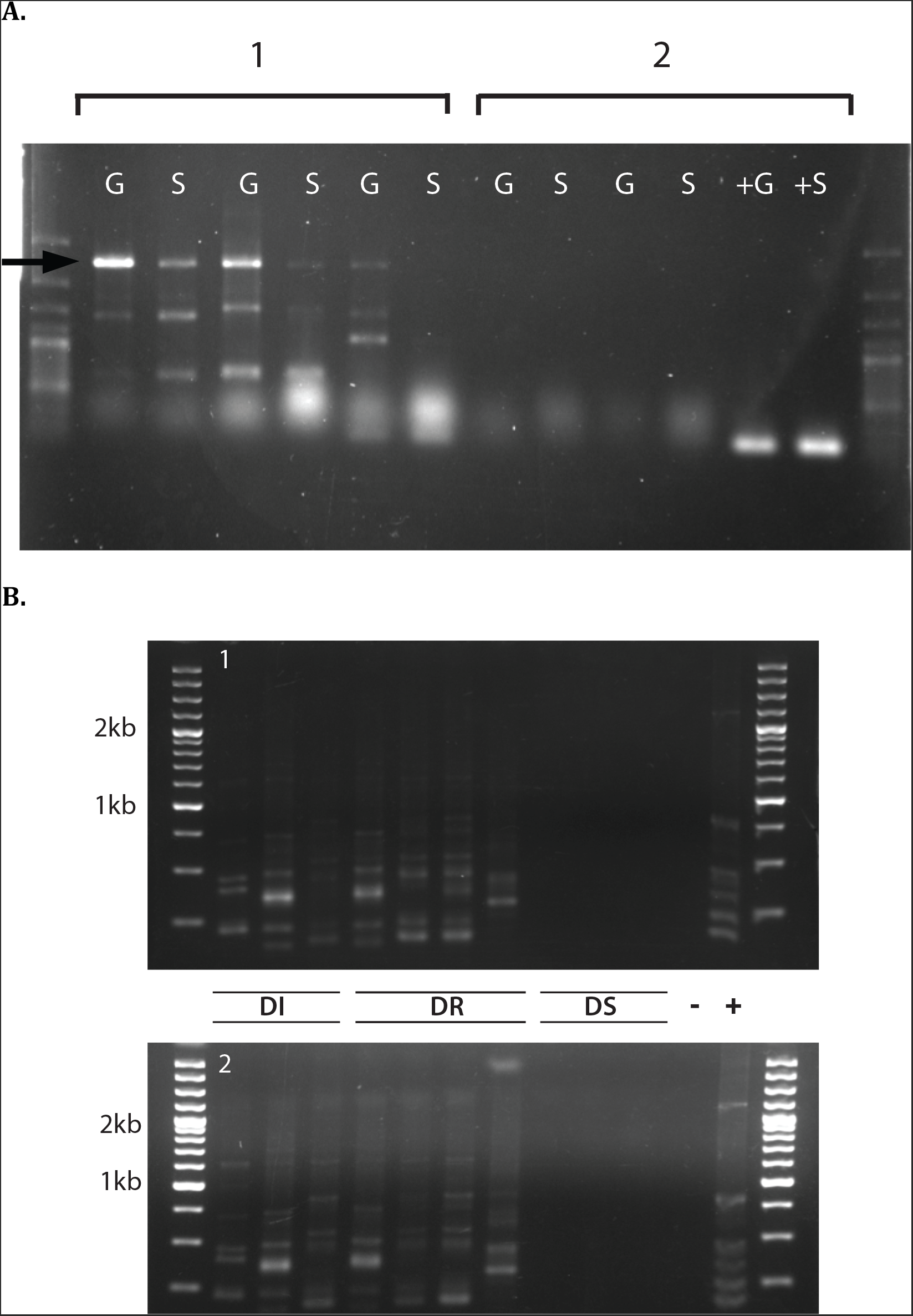
rtPCR for P-element in different line classifications. **A)** RT-PCR product from DI and DS lines. Group 1 consists of female flies from three DI lines roughly dissected into soma (S) and ovaries (G). The top band, at 2.6kb, corresponds to the expected size of the full-length P-element transcript after splicing. Group 2 consists of female flies from two DS lines with a positive control (*D. melanogaster* Harwich strain). **B)** PCR product from DI, DR & DS lines following reverse transcription with either forward (1) or reverse (2) primers.

**S1 Table. Fly strains used in this study.** Table includes time and date of collection, and collector. The fraction of strains with P-element are also shown for each collection.

**S2 Table. Association between cytotypes of *D. simulans* strains and infection status for individual TE families.** Number of reads mapping shows the number of reads which mapped to the TE in the expression data (SRA:PRJEB7936). The Z-value and P-value are from a binomial generalised linear model used to analyze the association between the TE copy number (found in the Florida sequence data, SRA:PRJEB7936, PRJNA308281) and the number of hybrid dysgenesis seen in an reciprocal crosses to an M line (see also S6 Fig); TEs causing the dysgenesis would be expected to show a significant relationship between dysgenesis in the cross where the Florida line is the male parent. Also shown is the number of DI strains and DS strains from which a TE could be amplified. Note that individual exons for the P-element could be amplified from all 39 DI strains; the results shown here are for amplification of the full-length P-element.

**S3 Table. Generalized linear model fit to the data on PCR presence/absence data.** The model used fit the year and region from which strains were collect (see S1 Table), plus an interaction term, to the presence or absence of P-element in that strain, and used a binomial error model. (I.e, the call in R glm(formula = cbind(present,absent) ∼ year ^*^ region, family = “binomial”)

**S4 Table. Primers used for identifying the presence or absence of P-element.**

